# Synergistic anticancer effect of plasma-activated infusion and salinomycin by targeting autophagy and mitochondrial morphology

**DOI:** 10.1101/2020.10.14.340208

**Authors:** Takashi Ando, Manami Suzuki-Karasaki, Miki Suzuki-Karasaki, Jiro Ichikawa, Toyoko Ochiai, Yukihiro Yoshida, Hirotaka Haro, Yoshihiro Suzuki-Karasaki

## Abstract

Non-thermal atmospheric plasma (NTAP)-activated liquids have emerged as new promising anticancer agents because they preferentially injure malignant cells. Here, we report plasma-activated infusion (PAI) as a novel NTAP-based anti-neoplastic agent. PAI was prepared by irradiating helium NTAP to form a clinically approved infusion fluid. PAI dose-dependently killed malignant melanoma and osteosarcoma cell lines showed much lower cytotoxic effects on dermal and lung fibroblasts. We found that PAI and salinomycin (Sal), an emerging anti-cancer stem cell agent, mutually operated as adjuvants. Combined administration of PAI and Sal was much more effective than single agent application in reducing the growth and lung metastasis of osteosarcoma allografts with minimal adverse effects. Mechanistically, PAI explicitly induced necroptosis and increased the phosphorylation of receptor-interacting protein 1/3 in a rapid and transient manner. PAI also suppressed the ambient autophagic flux by activating the mammalian target of rapamycin pathway. PAI increased the phosphorylation of Raptor, Rictor, and p70-S6 kinase, along with decreased LC3-I/II expression. In contrast, Sal promoted autophagy. Moreover, Sal exacerbated the mitochondrial network collapse caused by PAI, resulting in aberrant clustering of fragmented mitochondrial in a tumor-specific manner. Our findings suggest that combined administration of PAI and Sal is a promising approach for treating these apoptosis-resistant cancers.

## 1. INTRODUCTION

Osteosarcomas (OS) are malignant bone tumors accompanied by tumorous cartilage/bone or osteoid matrix formation. These tumors account for 15–20% of all malignant bone tumors, accounting for the highest percentage of all tumors (Morrow et al., 2018; Siegel et al., 2015). The incidence of OS is relatively low at 4.8 new cases per 1,000,000 individuals; however, most patients are children/adolescents aged <20 years. OS stages are classified as stage I (low-level malignancy) and stage II (high-level malignancy) according to the histological grade. Standard treatment for stage I tumors is surgery alone. Stage II or higher-grade tumors are therapeutically problematic. Standard treatment for stage IIA, IIB, or III tumors is a multidisciplinary approach consisting of pre- and post-operative chemotherapy and surgery. The 3-year overall survival rate ranges from 61% to 82%. For OS, combination chemotherapy with methotrexate, doxorubicin, and cisplatin is primarily used. However, 35–45% of patients with OS are insensitive to chemotherapy, and their 5-year survival rate is only 5–20% (Luetke et al., 2014). No other new drug with a high response rate has been developed and it remains difficult to improve prognosis using conventional chemotherapeutic regimens. Thus, an innovative therapy for overcoming drug resistance in OS is urgently required.

Non-thermal atmospheric plasma (NTAP) injures a variety of tumor cell types while sparing their non-transformed counterparts under optimal conditions (Keidar et al., 2011; Zucker et al., 2012; Guerrero-Preston et al., 2014; Ishaq et al., 2014). These properties have attracted much attention in cancer therapy. As with standard direct NTAP treatment, plasma-activated liquids (PALs) such as NTAP-irradiated media, solutions, and buffers exhibit strong anticancer activity (Girard et al., 2016; Van Boxem et al., 2017; Yan et al., 2018). Of these, plasma-activated medium (PAM) has been the most widely studied. PAM is conventionally produced by subjecting culture media such as Dulbecco’s modified Eagle’s medium (DMEM) to NTAP irradiation. Similarly to NTAP, PAM has a high potential to kill cancer cells including malignant melanoma (MM) and OS cells and evidence suggests that it acts in a tumor-specific manner (Saito et al., 2016; Tornin et al., 2019; Azzariti et al., 2019). However, conventional PAM includes many media components, such as the pH indicator phenol red. This compound undergoes specific metabolism to form a toxic substance in the liver (Driscoll et al., 1982). Thus, PAM may damage the liver in patients. As an alternative and safer NTAP-based tool, we developed a plasma-activated infusion (PAI) by irradiating a clinically approved infusion fluid with NTAP. As the infusion fluid consists of glucose, lactate, sodium chloride, and potassium chloride, the resulting PAI potentially contains no toxic substances and is expected to be safe for clinical use.

Autophagy is a primary catabolic process in which cellular components and damaged organelles undergo degradation. Three different types of autophagy have been identified: macro autophagy (referred hereafter as autophagy), micro autophagy [autophagy of organelles such as mitochondria and endoplasmic reticulum (ER)], and chaperone-mediated autophagy. The process of autophagy is complicated and includes induction of a phagophore in the cytoplasm, its elongation and autophagosome formation, the fusion of the autophagosome with lysosomes, and degradation of autophagosomal contents (Codogno and Meijer, 2005; Maiuri et al., 2007; Mizushima et al., 2011). These events are strictly controlled by autophagy-related genes (Codogno and Meijer, 2005). Autophagy copes with cellular stresses such as starvation and supplies energy and metabolic precursors. It is negatively regulated by the mammalian target of rapamycin complex 1/2 (mTORC1/2) in response to insulin and amino acid signals. During nutrient deprivation, this negative regulation by mTORC1/2 is alleviated, resulting in autophagy induction (Codogno and Meijer, 2005; Diaz-Troya et al., 2008; Dennis et al., 2011). Autophagy also contributes to removing damaged organelles such as mitochondria and ER via mitophagy and ERphagy, respectively. Accordingly, autophagy is essential for the survival of cancer cells by satisfying their high-energy demands and by removing damaged organelles (Ouyang et al., 2012; Bhutia et al., 2013). However, intensive and persistent activation of autophagy leads to programmed cell death, known as autophagic cell death (Gozuacik et al., 2004; Fulda et al., 2015; Fulda, 2017).

Salinomycin (Sal) is a naturally occurring polyether antibiotic that acts on potassium and calcium ionophores and has been used as an anticoccidial agent in the poultry industry for a long time. Sal has recently been considered as a promising anticancer drug because of its capacity to selectively kill cancer stem cells and multidrug-resistant cancer cells (Gupta et al., 2009; Dewangan et al., 2017; Kaushik et al., 2018). Several reports have demonstrated the participation of autophagic cell death in the anticancer effect of Sal (Verdoodt et al., 2012; Jangamreddy et al., 2013; Li et al., 2013; Mirkheshti et al., 2016). In contrast, inhibition of autophagy and induction of apoptosis plays a critical role in the anticancer effect of Sal (Endo et al., 2017; Pellegrini et al., 2016; Yue et al., 2013; Klose et al., 2014).

Different anticancer drugs, including temozolomide, epirubicin, and sorafenib, induce autophagy which contributes to drug resistance in various cancer cell types (Kanzawa et al., 2004; Knizhnik et al., 2013; Guo et al., 2016; Prieto-Dominguez et al., 2016). Autophagy also contributes to resistance to tumor necrosis factor-related apoptosis-inducing ligand (TRAIL) in colorectal cancers and hepatoma (He et al., 2012; Knoll et al., 2016; Lim et al., 2016). We previously demonstrated that in OS and MM cells, TRAIL induces robust autophagic flux whose suppression increases TRAIL-induced apoptosis (Ito et al., 2018; Onoe-Takahashi et al., 2019). Together, autophagy suppression represents a promising approach for overcoming drug resistance in cancer cells. Our preliminary data indicate that PAI can injury TRAIL-resistant OS and MM cells. Moreover, PAI can mimic the biological activities of simultaneous administration of TRAIL and autophagy inhibitors. We predicted that PAI can elicit anticancer activity by compromising autophagy. Additionally, if Sal can modulate autophagy, PAI and Sal may function cooperatively. This study was conducted to test these hypotheses in MM and OS cells.

## 2. MATERIALS AND METHODS

### 2.1 Materials

All chemicals were purchased from Sigma Aldrich (St. Louis, MO, USA) unless otherwise specified. Soluble recombinant human TRAIL was obtained from Enzo Life Sciences (Farmingdale, NY, USA). The pan-caspase inhibitor Z-VAD-FMK was purchased from Merck Millipore (Darmstadt, Germany). All insoluble reagents were dissolved in dimethyl sulfoxide and diluted with high glucose-containing DMEM supplemented with 10% fetal bovine serum (FBS) or Hank’s balanced salt solution (pH 7.4, Nissui Pharmaceutical Co., Ltd., Tokyo, Japan) (final dimethyl sulfoxide concentration, <0.1%) prior to use.

### 2.2 Cell Culture

The human melanoma cell line A2058 (cell number IFO 50276) was obtained from the Japanese Collection of Research Bioresources (JCRB) Cell Bank of National Institutes of Biomedical Innovation, Health, and Nutrition (Osaka, Japan). Human fetal lung fibroblasts, WI-38 (cell number JCRB9017), were obtained from JCRB. Human dermal fibroblasts from the facial dermis were obtained from Cell Applications (San Diego, CA, USA). Human osteosarcoma HOS (RCB0992), MG63 (RCB1890), Saos-2 (RCB0428), 143B (RCB0701), and murine osteosarcoma LM8 (RCB1450) cells were purchased from Riken Cell Bank (Tsukuba, Japan). The cells were maintained in 10% FBS (GIBCO^®^, Life Technologies, Carlsbad, CA, USA) containing DMEM (GIBCO^®^, Life Technologies) (FBS/DMEM) supplemented with, 100 U/mL penicillin and 100 μg/mL streptomycin at 37°C in a humidified atmosphere with 5% CO_2_.

### 2.3 PAI Preparation

NTAP was generated from helium using a PCT-DFJIM-02 model damage free multi-gas plasma jet (Plasma Concept Tokyo, Tokyo, Japan), which generates a capacitively coupled plasma. The typical frequency was 20 kHz, with a peak voltage of 1 kV, current of 30 mA, and a helium flow rate of 3 L/min. PAI (1 mL) was made by irradiating plasma from above at a distance of 20 mm to 5 mL of Soldem 3A (TERUMO, Tokyo, Japan) for 1 or 5 min. The original PAI was diluted to a final concentration of 6.3–50% with 10% FBS/DMEM (for cell experiments) or Hank’s balanced salt solution (for biochemical experiments) and was indicated as PAI (6.3–50%).

### 2.4 Quantitation of H_2_O_2_ in PAI

The concentration of H_2_O_2_ in PAI was measured using the Amplex Red Hydrogen Peroxide/Peroxidase Assay Kit (Thermo Fisher Scientific, Waltham, MA, USA) according to the manufacturer’s protocols. This assay is based on the formation of the oxidant product resorufin by the reaction of the Amplex Red reagent (10-acetyl-3, 7-dihydroxyphenoxazine) with H_2_O_2_ in a 1:1 stoichiometry in the presence of peroxidase. Briefly, samples were diluted to 20-fold and placed on a 96-well plate (50 μL/well). Next, 50 μL of a working solution of 100 μM of Amplex Red reagent and 0.2 U/mL of horseradish peroxidase was added to the well and incubated at room temperature for 30 min. Absorbance at 570 nm was measured using a microplate reader (Nivo 3F Multimode Plate Reader, PerkinElmer, Waltham, MA, USA). The concentrations of H_2_O_2_ were calculated using a standard curve, prepared using authentic H_2_O_2_ included with the kit.

### 2.5 Measurements of Intracellular ROS Generation

LM8 OS cells were cultured in 6-well plates for 24 h and then exposed to PAI (50%) or H_2_O_2_ (100 μM) for 1 h. Cells were labeled with DCFH-DA FITC antibody using the Reactive Oxygen Species (ROS) Detection Assay Kit obtained from BioVision, Inc. (Milpitas, CA, USA) according to the manufacturer's instructions. Data were collected using a FACSCalibur (BD Biosciences, Franklin Lakes, NJ, USA) and were analyzed using CellQuest Pro (BD Biosciences) and FlowJo software (TreeStar, Ashland, OR, USA). Images of ROS formation in LM8 live cells in each well were taken with the Fluorescence Microscope FLUOVIEW FV10i (Olympus, Tokyo, Japan) (Ex/Em 495/529 nm). Experiments were performed in triplicate.

### 2.6 Animals

Homozygous wild-type (WT) C3H/HeJJcl mice were purchased from CLEA Japan (Tokyo, Japan). The mice were housed at 22–24°C with a 12-h light/dark cycle with standard mouse chow and water provided *ad libitum*. All experiments with mice were conducted according to the Guidelines for Proper Conduct of Animal Experiments, Science Council of Japan, and the protocols were approved by the Animal Care and Use Committee (No. 17–11) at the University of Yamanashi.

### 2.7 Cell Viability Assay

Cell viability was measured by the WST-8 assay using Cell Counting Reagent SF (Nacalai Tesque, Inc., Kyoto, Japan) or Cell Counting Kit-8 (Dojindo Molecular Technologies, Inc., Kumamoto, Japan) according to the manufacturer's instructions. These methods are colorimetric assays based on the formation of a water-soluble formazan product (WST-8). Briefly, cells were seeded at a density of 8 × 10^3^ cells/well in 96-well plates (Corning, Inc., Corning, NY, USA) and cultured with agents to be tested for 72 h at 37°C before adding 10 μL cell counting reagent and further incubation for 2 h. Absorbance at 450 nm was measured using a microplate reader (Nivo 3F, PerkinElmer or SH-1100R (Lab), Corona Electric Co., Ltd., Ibaraki, Japan).

### 2.8 Cell Death Assay

Cells were cultured in 6-well plates for 24 h and then exposed to several concentrations of PAI for 24 to 48 h. To evaluate apoptotic cell death, the cells were retrieved using Versene (GIBCO^®^, Life Technologies) and incubated with Annexin V and 7AAD (BD Biosciences) for 15 min. Data were collected using a FACSCalibur (BD Biosciences). Data obtained were analyzed by CellQuest Pro (BD Biosciences) and FlowJo software (TreeStar). Experiments were performed in triplicate. Overall cell death was evaluated using a commercially available kit (Live/Dead Viability/Cytotoxicity Kit; Invitrogen, Carlsbad, CA, USA) according to the manufacturer’s instructions as described previously (Pellegrini et al., 2016). Briefly, the cells were cultured on an 8-well imaging chamber (Imaging Chamber 8 CG, Zell-Kontakt GmbH, Nörten-Hardenberg, Germany) and treated with the agents to be tested for 24 h at 37°C. The cells were stained with 4 μM each of calcein-AM and ethidium bromide homodimer-1 (EthD-1) to label live cells in green and dead cells in red, respectively. Images were obtained using a BZ X-710 Digital Biological Microscope (Keyence Corporation, Osaka, Japan) equipped with a 40×, 0.60 numerical aperture (NA) LUCPlanFL N objective (Olympus) and were analyzed using BZ-H3A application software (Keyence Corporation).

### 2.9 Tumor Growth and Metastasis

The ability of PAI or Sal to reduce tumor growth *in vivo* was evaluated using allograft transplants of LM8 murine OS cells in mice. Male C3H/HeJJcl mice of 8 weeks age were administered general anesthesia with isoflurane (ISOFLU^®^) (Abbott Laboratories, North Chicago, IL, USA) and oxygen. LM8 murine osteosarcoma cells (2 × 10^6^ cells/mouse) in 0.1 mL DMEM were injected subcutaneously into the back of the mice on day 0. Three times per week after day 7, 200 μL of PAI (50%) and Sal (3 mg/kg) alone or in combination was administered intravenously to 6 mice in each group. The mice were weighed, and the size of the primary tumors was measured weekly. On day 35, all mice were sacrificed (Supplementary Figure 6A). Tumors and lungs of mice were fixed in 10% formalin neutral buffer solution for 3 days. Tumors and lungs were paraffin-embedded and consecutive 5-μm sections were stained with hematoxylin-eosin purchased from Merck.

### 2.10 Immunohistochemical (IHC) Staining

IHC staining was performed as previously described (Sato et al., 2016). IHC staining with primary antibodies against LC3-I/II (D3U4C) (#12741, 1:2000), Beclin-1 (D40C5) (#3495, 1:400) (Cell Signaling Technology, Danvers, MA, USA) and Ki67 (SP6) (ab16667, 1:100) (Abcam, Cambridge, UK) was performed using the Dako Liquid DAB+Substrate Chromogen System (Glostrup, Denmark) according to the manufacturer's specifications, followed by counterstaining with hematoxylin.

### 2.11 Autophagy Assay

Cells were cultured in 6-well plates for 24 h and then exposed to several concentrations of PAI for 24–48 h. The cells were stained and analyzed using the Cyto-ID^®^ Autophagy Detection Kit obtained from Enzo Life Sciences according to the manufacturer's instructions. Data were collected using a FACSCalibur (BD Biosciences) and analyzed using CellQuest Pro (BD Biosciences) and FlowJo software (TreeStar). Experiments were performed in triplicate.

### 2.12 Western Blotting Analysis

Cells were washed in Ca2^+^-, Mg2^+^-free PBS, lysed during 15 min shaking in CelLytic M lysis buffer (Merck KGaA) with a protease inhibitor cocktail and phosphatase inhibitor cocktail (both from Sigma). Protein concentrations were determined using a BCA protein assay (Thermo Fisher Scientific) according to the manufacturer’s protocol. Equal amounts of protein from each sample were analyzed as previously described by immunoblotting with primary antibodies against cleaved caspase-3 (Asp175) (5A1E) (#9664, 1:1000), cleaved caspase-8 (Asp387) (D5B2) (#8592, 1:1000), phospho-RIP1 (Ser166) (#31122, 1:1000), RIP (D94C12) (#3493, 1:1000), phospho-P70-S6 (Thr389) (108D2) (#9234, 1:1000), P70-S6 (49D7) (#2708, 1:1000), phospho-Rictor (Thr1135) (D30A3) (#3806, 1:1000), Rictor (53A2) (#2114, 1:1000), phospho-Raptor (Ser792) (#2083, 1:1000), LC3-I/II (D3U4C) (#12741, 1:1000), and GAPDH (D16H11) (#5174, 1:1000) obtained from Cell Signaling Technology; phospho-RIP3 (phospho S232) (EPR9516(N)-25) (ab195117, 1:1000), Raptor (EP539Y) (ab40768, 1:500) obtained from Abcam; and phospho-P62 (Ser351) (SQSTM1) (PM074, 1:500) and P62 (SQSTM1) (PM045, 1:1000) obtained from MBLI (Woburn, MA, USA), were used. Images were captured using a LAS-4000 camera system from Fujifilm (Tokyo, Japan).

### 2.13 Live-Cell Mitochondrial Network Imaging

The mitochondrial network in live cells was analyzed as previously described (Saito et al., 2016). Briefly, cells in FBS/DMEM (3 × 10^4^/well) adherent on 8-well chambered coverslips were treated with the agents to be tested for 24 h at 37°C in a 5% CO_2_ incubator. After removing the medium by aspiration, the cells were washed with fresh FBS/DMEM and stained with 20 nM MitoTracker Red CMXRos for 1 h at 37°C in the dark in a 5% CO_2_ incubator. In some experiments, the nuclei were counter-stained with 1 mg/mL of Hoechst 33342. The cells were then washed with and immersed in FluoroBriteTM DMEM (Thermo Fisher Scientific). Images were obtained using a BZ X-710 Digital Biological Microscope (Keyence) equipped with a 100 ×, 1.40 n.a. UPlanSApo Super-Apochromat, coverslip-corrected oil objective (Olympus). Images were analyzed using BZ-H3A application software (Keyence) and ImageJ software (NIH, Bethesda, MD, USA). For each sample, the morphology of mitochondria in 30–60 cells was analyzed, and the percentages of four different types, i.e., tubular/fused, fragmented, and swollen/clustered, were calculated.

### 2.14 Statistical Analysis

Data are presented as mean ± standard deviation (SD) or standard error (SE) and were analyzed by one-way analysis of variance followed by Tukey’s post hoc test using add-in software with Excel 2016 for Windows (SSRI, Tokyo, Japan). For some experiments, significance was determined using Student’s *t*-test after an F-test. *P* < 0.05 was considered as statistically significant.

## 3. RESULTS

### 3.1 PAI Contains Oxidants and Evokes the Subsequent Intracellular ROS Generation

Various reactive oxygen/nitrogen species (ROS/RNS) are generated in the air and liquid phases following NTAP irradiation and relatively long-lived species are thought to be responsible for the anticancer activity (Morrow et al., 2018; Zucker et al., 2012; Guerrero-Preston et al., 2014). Of these, H_2_O_2_ is found in various types of PALs and was shown to be the primary mediator of their anticancer activity (Keidar et al., 2011; Zucker et al., 2012; Guerrero-Preston et al., 2014; Ishaq et al., 2014; Girard et al., 2016). Therefore, we first quantitated the amounts of H_2_O_2_ in PAI. We previously found that the anticancer activity of PAM was correlated with the time of NTAP irradiation and inversely correlated with the volume of the target solution (Saito et al., 2016). Accordingly, the ratio of time and volume is critical for determining efficacy. As expected, PAI contained substantial amounts of H_2_O_2_, which increased as the ratio increased (Figure 1A); the amounts were 8.37 ± 0.29 and 124.6 ± 27.1 (μM) (n = 3) for NTAP irradiation of 12 s (0.2 min/mL) and 60 s(1 min/mL), respectively. Previously, we demonstrated that PAM stimulates intracellular ROS generation, including that of the mitochondrial superoxide (Saito et al., 2016). Flow cytometric analysis showed that like PAM, PAI treatment increased the intracellular ROS level in LM8 OS cells (Figure 1B). We also detected robust ROS generation in live adherent cells (Figure 1C). In this case, a much stronger ROS signal was observed with PAI than with H_2_O_2_ (100 μM). These results indicate that PAI contains H_2_O_2_ and can evoke subsequent intracellular ROS generation.

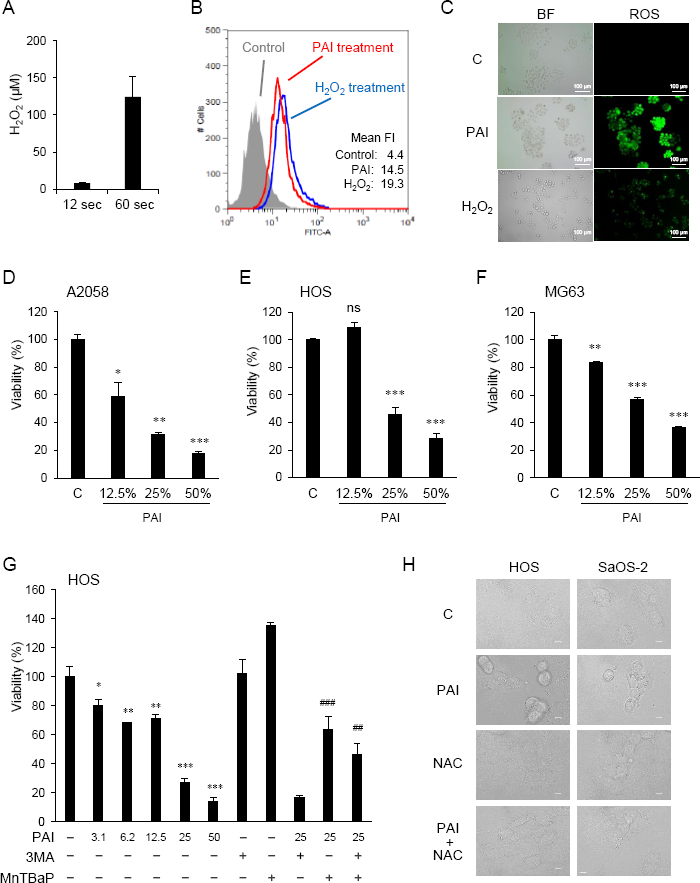

### 3.2 PAI Reduces Cell Viability in MM and OS cells in an ROS-Dependent Manner

We examined the ability of PAI to reduce cell viability. Cells were treated with varying concentrations of PAI for 72 h and analyzed for their viability in a WST-8 cell growth assay. PAI treatment resulted in a dose-dependent decrease in the viability of A2058 MM, HOS, and MG63 OS cells (Figure 1D F) compared to control cells treated with the vehicle (infusion fluid). PAI was also effective in other OS cell lines including LM8 and Saos-2 (Figure 2) and 143B cells (Figure 4). As we previously reported that ROS (Saito et al., 2016) and autophagic cell death (Ito et al., 2018) mediated the anticancer effect of PAM, we examined the possible roles of these events in the effects of PAI. The superoxide dismutase mimetic MnTBaP significantly inhibited cell death, whereas the autophagy inhibitor 3-MA did not (Figure 1G). Moreover, the broad-spectrum antioxidant *N*-acetylcysteine completely prevented PAI induced cell damage in HOS and Saos-2 cells (Figure 1H). These results suggest that ROS, but not autophagy, plays a role in mediating cell death induced by PAI.

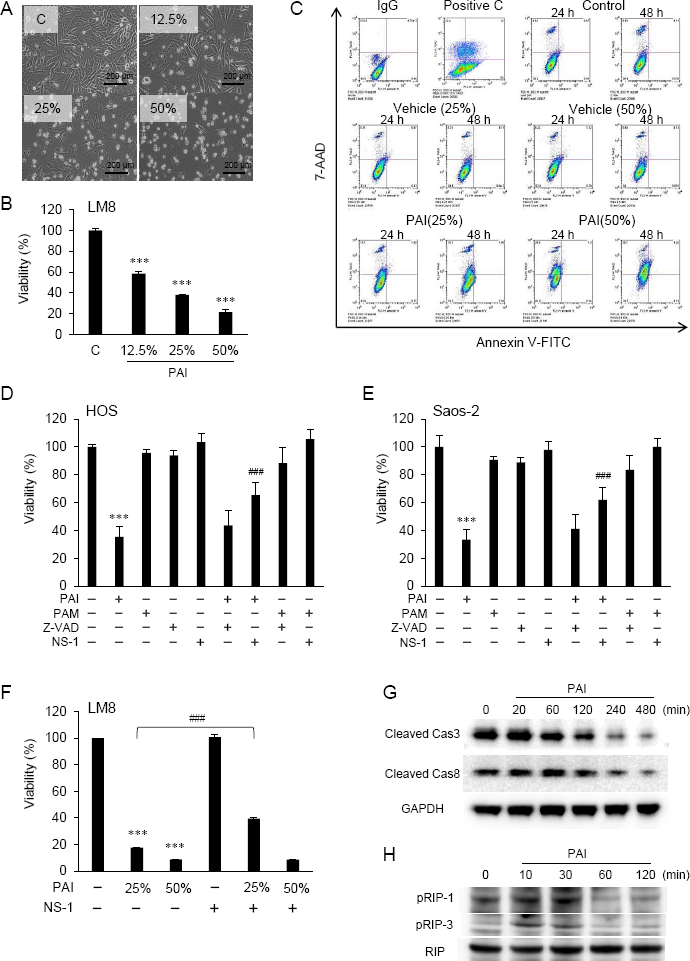

### 3.3 PAI Induces Necroptosis

To determine the cell death modality, we observed the morphology of cells following PAI treatment under a light microscope. Figure 2A shows the results obtained in LM8 cells. The adherent spindle cells were converted to less adherent, shrunken, and round cells in a dose-dependent manner. These morphological changes were associated with a dose-dependent decrease in cell viability (Figure 2B). Double staining with FITC-conjugated Annexin V and 7-AAD followed by flow cytometric analyses showed a dose- and time-dependent increase in Annexin V-negative, 7-AAD-positive cells and minimal increase in Annexin V-positive cells with PAI treatment (Figure 2C). These results suggest that PAI preliminarily causes necrotic cell death and that apoptosis plays a minor role. To confirm this, we examined the effect of pharmacological inhibitors specific for apoptosis and necroptosis. Receptor interacting protein (RIP) kinases 1 and 3 have essential scaffolding functions that activate necroptosis in cells. In HOS and Saos-2 cells, necrostatin-1 (NS-1), a specific inhibitor of RIP1 kinase (RIPK), inhibited cell death significantly, whereas the broad-spectrum caspase inhibitor Z-VAD-FMK (ZVAD) had minimal effects (Figure 2D, E). Additionally, NS-1 significantly reduced PAI-induced cell death in LM8 cells treated with PAI (25%) but not PAI (50%) (Figure 2F). Next, we examined the status of caspase-3, caspase-8, RIPK1, and RIPK3 activity in the cells. Western blot analysis showed that the expression of activated caspase-3 and caspase-8 (cleaved caspase-3 and caspase-8) were decreased following PAI treatment (Figure 2G, Supplementary Figure 1A, B). In contrast, expression of activated RIPK1 and RIPK3 (phosphorylated-RIPK1 and -RIPK3) was increased (Figure 2H, Supplementary Figure 2A, B). This effect was rapid and transient; the increases were initially observed within 10 min, reached a maximum after that, and declined to the basal levels by 60 min. These results indicate that PAI primarily induces necroptosis in OS cells.

### 3.4 PAI Reduces Autophagic Flux by Promoting the mTORC Pathway

The results shown in Figure 1 suggest that autophagic cell death plays a minor role in the anticancer effect of PAI. In cancers, autophagy exerts both cytocidal and cytoprotective roles. In support of this view, we previously reported that human MM and OS cells exhibit a substantial level of autophagic flux even under stress-free and nutritional conditions, and that ambient autophagy prevented the cells from spontaneous and TRAIL-induced apoptosis (Ito et al., 2018; Onoe-Takahashi et al., 2019). Therefore, next, we determined the possible role of cytoprotective autophagy. We investigated the effect of PAI on the autophagic flux in OS cells. When Cyto-ID, the specific probe of autophagosome formation, was analyzed by flow cytometry, rapamycin treatment substantially increased the signal, whereas PAI significantly reduced it (Figure 3A). mTORC plays a pivotal role in regulating autophagy by inhibiting phagophore formation (Codogno and Meijer, 2005; Diaz-Troya et al., 2008; Dennis et al., 2011). To determine whether this complex participates in the suppression of autophagy, we examined the effect of PAI on mTORC activation by western blot analysis. The results showed that PAI increased the phosphorylation of p70-S6 kinase, the substrate of mTOR kinase1/2, as well the phosphorylation of Rictor and Raptor, which are components of these two kinases. PAI decreased the protein levels of LC3-I/II, an autophagosomal marker, over time (Figure 3B, Supplementary Figure 3, 4).

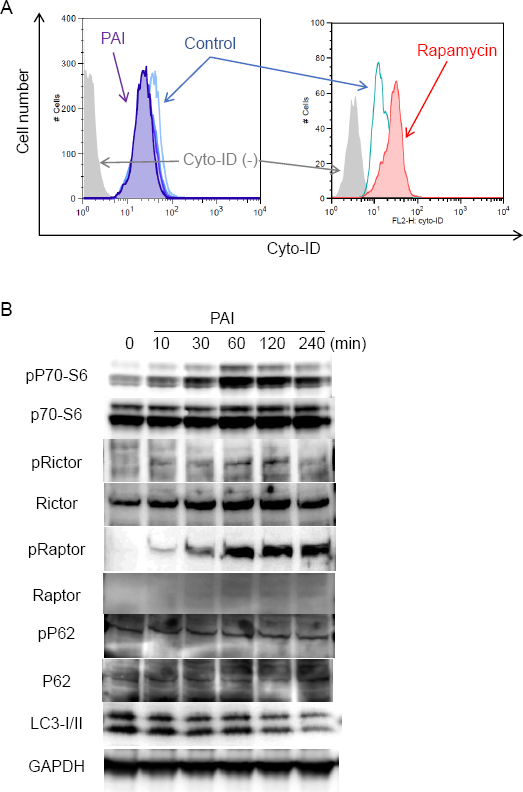

### 3.5 PAI and Sal Mutually Act as an Adjuvant in a Tumor-specific Manner

Although the mechanisms by which Sal exhibited the anticancer effect remain unknown, recent studies demonstrated that Sal targets autophagy (Dewangan et al., 2017; Kaushik et al., 2018; Verdoodt et al., 2012; Jangamreddy et al., 2013; Li et al., 2013; Mirkheshti et al., 2016; Endo et al., 2017; Pellegrini et al., 2016; Yue et al., 2013; Klose et al., 2014). Therefore, we determined the effect of Sal on autophagy flux. Western blot analysis showed that Sal treatment resulted in increased phosphorylation of p62 (phospho-p62) and LC3-I/II in LM8 cells (Figure 4A, Supplementary Figure 5). These increases were observed as rapidly as 30 min following Sal treatment and became clearer over time up to 120 min. Thus, Sal promotes apoptosis in cells. Next, we examined whether Sal could induce cell death in our cell systems. As shown in Figure 4B, Sal induced cell death, which was characterized by an increase in the number of detached and round cells along with membrane blebbing. Thus, the effect resembled that caused by PAI. Sal at concentrations of ≥3 μM significantly reduced cell viability, and Z-VAD-FMK minimally inhibited this effect while Z-VAD-FMK plus NS-1 partially blocked it (Figure 4C). Combined application of PAI and Sal appeared to cause more severe cell damage than either agent alone. To confirm this cooperative action, we examined the combined effect of PAI and Sal treatment on cell viability. We found that 3 μM was a borderline range in which Sal had substantial or modest cytotoxic effect depending on cell types. When used with PAI, a greater extent of cell death was reproducibly observed compared with that caused by either single agent (Figure 4D). This effect was observed over a wide range of PAI (6.3–50%) in different OS cell lines (Figure 4E, F). In contrast, treatment with PAI (12.5–50%) for 72 h only modestly decreased the viability of WI-38 cells (maximum of 24% at 50%) (Figure 4G). Staining with calcein-AM and EthD-1 showed that treatment with PAI and Sal alone or in combination for 18 h substantially increased the proportion of EthD-1-positive (dead/damaged) cells in HOS cells along with decreased calcein-positive (live) cells, whereas they minimally increased the proportion of PI-positive cells and decreased the proportion of calcein-positive cells in human dermal and WI-38 lung fibroblasts (Figure 4H).

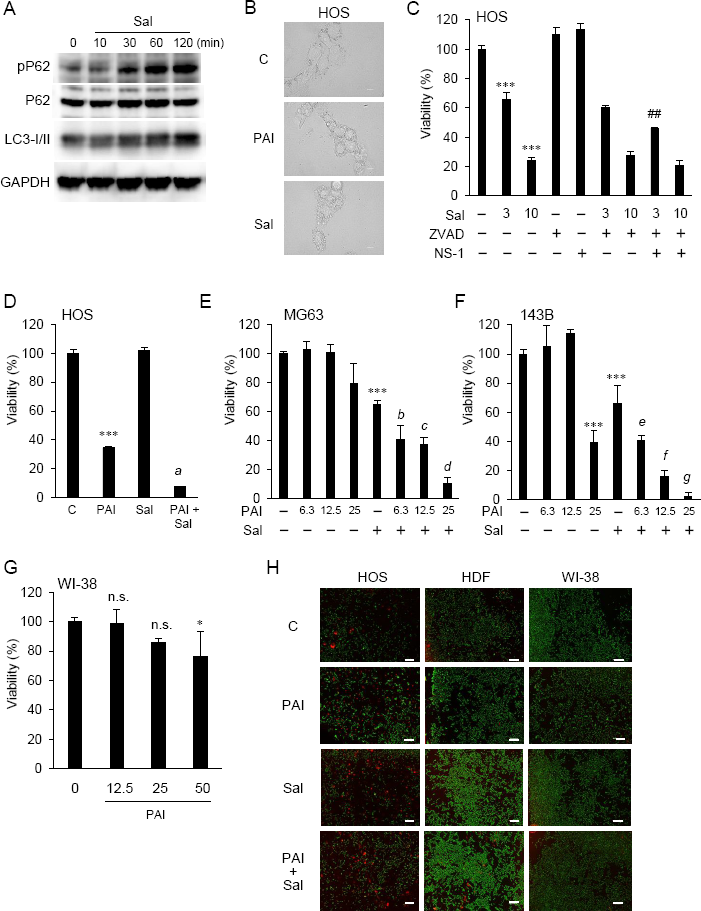

### 3.6 Combined Administration of PAI and Sal Reduces the Growth and Metastasis of OS Allografts with Minimal Adverse Effects

Next, we determined whether PAI and Sal operate cooperatively *in vivo*. LM8 can be transplanted into C3H mice to induce the development of tumors with high metastatic potential to the lung after inoculation into the skin. We assessed the anticancer activity of PAI and Sal alone or in combination in this allograft model (Supplementary Figure 6A). After subcutaneous inoculation of LM8 cells into mice, tumors exhibited rapid growth and reached a size of about 400 mm^3^ within 5 weeks. Treatment of the mice by intravenous administration of PAI and Sal 3 times weekly resulted in a significant (47.2% and 50.2%, respectively) reduction in the primary tumor size (Figure 5A, B, Supplementary Figure 6B). A more pronounced anticancer effect (24.9% reduction) was observed in mice treated by combined administration of PAI and Sal (Figure 5A, B). Additionally, histological analysis with hematoxylin-eosin stain revealed that PAI and Sal significantly decreased the colony number of metastatic nodules in the lungs of LM8-inoculated mice. Again, combined administration of PAI and Sal was more effective than single administration of either agent (Figure 5C, D). Despite the potent anticancer effect, there was a minimal difference in weight between the mice the treated and untreated groups and no obvious abnormality was observed in treated mice with this protocol (Figure 5E). These results indicate that combined administration of PAI and Sal has potent and tumor-selective anticancer effects *in vivo*. The impact of PAI and Sal on autophagy *in vivo* was determined by IHC analyses using specific antibodies against LC3-I/II and Beclin-1. LC3-I//II and Beclin-1 levels were lower in the PAI treatment group and higher in the Sal treatment group. These levels in the combined administration of PAI and Sal group were intermediate (Figure 6A–C). The tumor proliferation index marker Ki67 is intimately associated with tumor cell proliferation (Richardsen et al., 2017; Inwald et al., 2013). The Ki67 labeling index for the PAI or Sal administration group was significantly lower than that of the control group. Furthermore, combined treatment with these agents reduced this index more significantly than each monotherapy (Figure 6D, E). These results agree with our *in vitro* study.

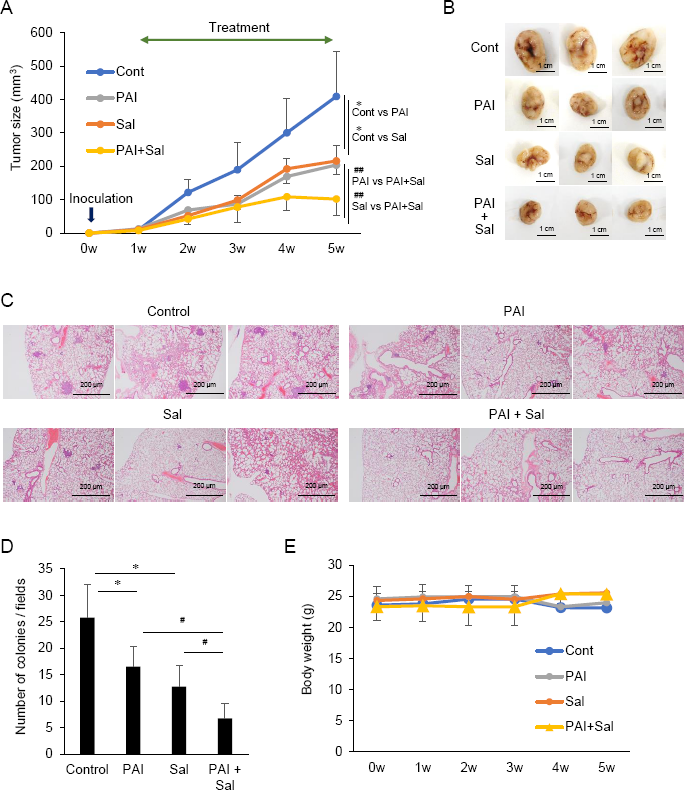

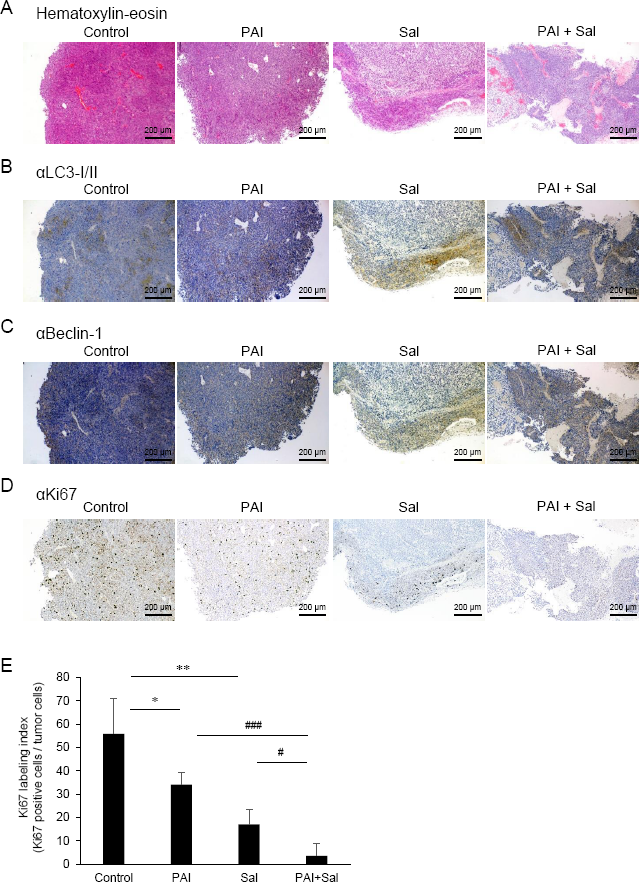

### 3.7 PAI and Sal Alter Mitochondrial Morphology Cooperatively

Previously, we demonstrated that PAM induced excessive mitochondrial fragmentation along with swelling and clustering in tumor cells. In contrast, mitochondrial morphology was minimally changed in non-transformed cells following PAM treatment (Saito et al., 2016). Therefore, we examined whether PAI also affects mitochondrial morphology in cancer cells. We stained the mitochondria in live cells with MitoTracker Red and observed them under a biological microscope. The nuclei were counter-stained with Hoechst 33342. Control cells were highly adherent and spindle cells, in which most mitochondria displayed a reticular network around the intact nuclei (Figure 7A, left panels). PAI dose-dependently altered mitochondrial morphology. Upon treatment with PAI (12.5%), the mitochondria were fragmented and some became swollen (Figure 7A, 2nd panels from left, 7B). At higher PAI concentrations (≥25%), most mitochondria became fragmented, swollen, and clustered at the perinuclear regions of one side of the nuclei (Figure 7A, 3rd and 4th panels from left, 7B). These changes were pronounced in less adherent, round, and blebbing cells. PAM was less effective than PAI in modifying mitochondrial morphology. PAM (25%) led to only modest mitochondrial fragmentation with little swelling (Figure 7A, right panels). We obtained similar results in A2058 cells (Figure 7C). In contrast, even the highest concentration of PAI (50%) led only to a modest mitochondrial fission in WI-38 fibroblasts with minimal swelling and clustering (Figure 7D). We also examined the possible impact of Sal on the mitochondrial morphology. Upon treatment with Sal (3 μM), the mitochondria underwent massive fragmentation, part of which was swollen in HOS cells (Figure 7E, 3rd panels from left, 7F). When administered together, PAI (25%) and Sal (1 μM) induced mitochondrial fragmentation with robust swelling and clustering in the cells (Figure 7D, right panels, 7F).

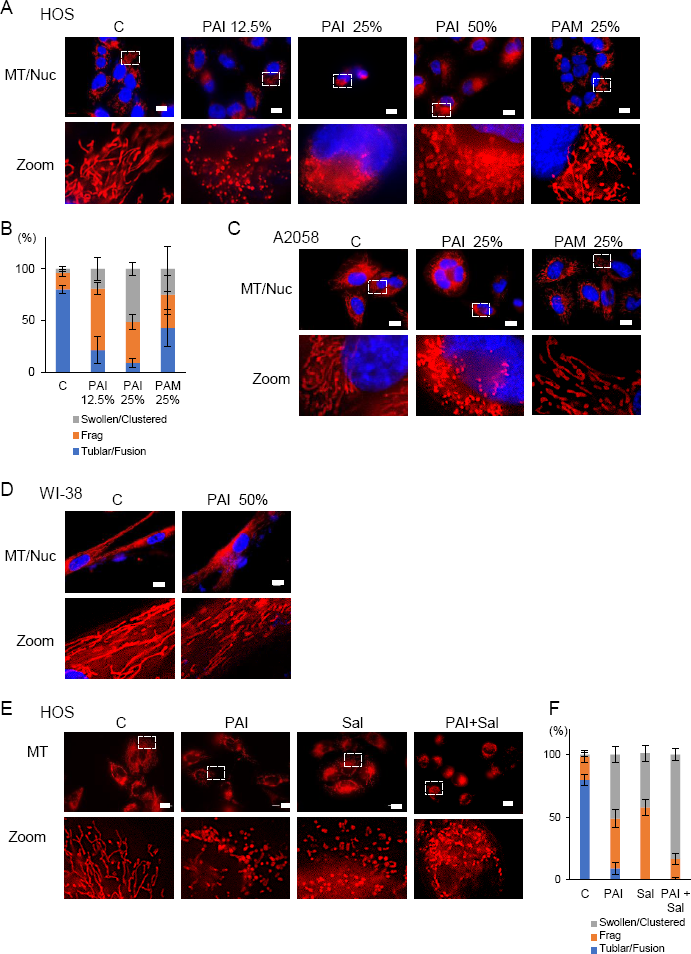

## 4. DISCUSSION

In the present study, we showed that PAI had potent anticancer activity against MM and OS cells by inducing cell death (Figure 1, 2, 4) and reduced OS tumor growth and metastasis to the lung *in vivo* (Figure 5, 6). Notably, PAI showed minimal cytotoxicity in non-transformed human dermal and lung fibroblasts (Figure 4), indicating that it preferentially injures tumor cells. PAI administration displayed no significant adverse effects including weight loss *in vivo* (Figure 5), supporting its tumor-selective action. The antioxidant MnTBaP and *N*-acetylcysteine significantly inhibited the anticancer effect of PAI (Figure 1), suggesting that like other PALs, PAI elicits the effect in a ROS-dependent manner. Consistent with this view, PAI contained H_2_O_2_ and increased the intracellular ROS (Figure 1). Our data indicated that unlike other PALs, including PAM, which mainly stimulate apoptosis (Girard et al., 2016; Van Boxem et al., 2017; Yan et al., 2018), PAI primarily increased necrotic cell death, which was minimally affected by caspase inhibition (Figure 2). In support of this, PAI triggered necroptotic signaling pathways by rapidly and transiently activating RIP1 and RIP3 kinases (Figure 2). Moreover, the necroptosis-specific inhibitor NS-1 significantly but not completely reduced cell death caused by PAI (Figure 2). These data suggest that PAI primarily targets necroptosis. This is consistent with the fact that ROS plays a critical role in regulating necroptosis (Fulda, 2016) and the cytotoxic effect of PAI (Figure 1, 2, 4). Although the reason for the limited effect of NS-1 is unknown, there are several possibilities. The simplest possibility is that another mode of programmed cell death mediates the effect of PAI. Alternatively, RIP1 and RIP3 kinases may be equally essential for transducing necroptotic cell death signaling. This idea is consistent with observations that PAI activated both kinases with similar kinetics (Figure 2). If both kinases were involved, it is logical that NS-1, which targets RIP1 signaling alone, failed to block the death signaling entirely. However, we did not observe complete inhibition of the anticancer effect of PAI even when expression of the RIP1 and RIP3 genes was downregulated using specific siRNAs (data not shown). Thus, inhibition of necroptosis may lead to the onset of another mode of programmed cell death. Further investigation is necessary to clarify the role of another cell death mechanism in the anticancer effect of PAI.

Aggressive tumors have varying cellular machinery that protects them from apoptosis caused by anticancer agents, thereby conferring drug-resistance. An emerging view is that apoptosis (type I cell death) is not the sole mode of programmed cell death, but several different modes of = occur in response to anticancer agents. Of these, autophagy (type II cell death), paraptosis (type III cell death), and necroptosis are known. Thus, recently, cancer therapy based on induction of non-apoptosis has been considered as an alternative approach to treat apoptosis-resistant cancer cells including MM and OS cells. We previously showed that PAM could induce autophagic cell death in MM and OS cells (Ito et al., 2018). PAM can increase the Cyto-ID signal and protein level of LC3-I/II and the autophagy inhibitor 3-MA and bafilomycin A1 can block cell death in these cells. The present study showed that PAI reduced the ambient Cyto-ID signal and LC3-I/II protein expression (Figure 3). Moreover, 3-MA minimally affected cell death caused by PAI (Figure 1). These data indicate that autophagic cell death plays a minor role in the anticancer effect. In support of autophagy suppression, PAI can activate mTORC1/2, the autophagy suppressive machinery; further, PAI increased the phosphorylation of Raptor, Rictor, and their substrate P70-S6 kinase (Figure 4). At present, the mechanisms by which PAI activates these pathways are unknown. However, ROS may play a role in this context because mTORC activation is regulated by oxidative stress through a redox-sensitive mechanism (Huang et al., 2002; Sarbossov et al., 2006; Reiling and Sabatini, 2006). Autophagy contributes to cancer cell survival and resistance to different types of anticancer drugs (Kanzawa et al., 2004; Knizhnik et al., 2013; Guo et al., 2016; Prieto-Dominguez et al., 2016; He et al., 2012; Knoll et al., 2016; Lim et al., 2016). Thus, the suppression of autophagy may also participate in the anticancer effect of PAI.

PAI and Sal mutually acted as an adjuvant *in vitro* (Figure 4). Notably, the combination reduced not only tumor growth but also metastasis to the lung while causing minimal adverse effects, including weight loss (Figure 5). The mechanisms underlying the cooperative actions between PAI and Sal are currently unknown. Notably, these two agents cooperatively disrupt the mitochondrial network. An emerging view is that mitochondrial network homeostasis plays a crucial role in regulating cancer cell survival, thereby serving as a promising target for cancer treatment (1-3). The delicate balance of mitochondrial fission and fusion is essential for maintaining functional mitochondrial morphology. Accordingly, excessive fission and fusion leads to mitochondrial dysfunction and cell death. Toxic concentrations of PAI explicitly caused excessive mitochondrial fragmentation, swelling, and clustering in MM and OS cells, whereas only a modest mitochondrial fission occurred in fibroblasts (Figure 7). Low concentrations of PAI primarily caused mitochondrial fragmentation with minimal swelling and clustering. We found that Sal showed similar effects. Interestingly, when PAI and Sal were administered together, substantial mitochondrial swelling and clustering occurred (Figure 7). Mitochondrial swelling is a hallmark of the opening of multiple mitochondrial permeability transition pores and mitochondrial clustering represents an irreversible denature of the mitochondria, which may compromise the mitochondrial network homeostasis. Therefore, swelling and clustering of fragmented mitochondria are critical in cell death and that exacerbation of these events may play a role in the cooperative actions between PAI and Sal (Figure 8).

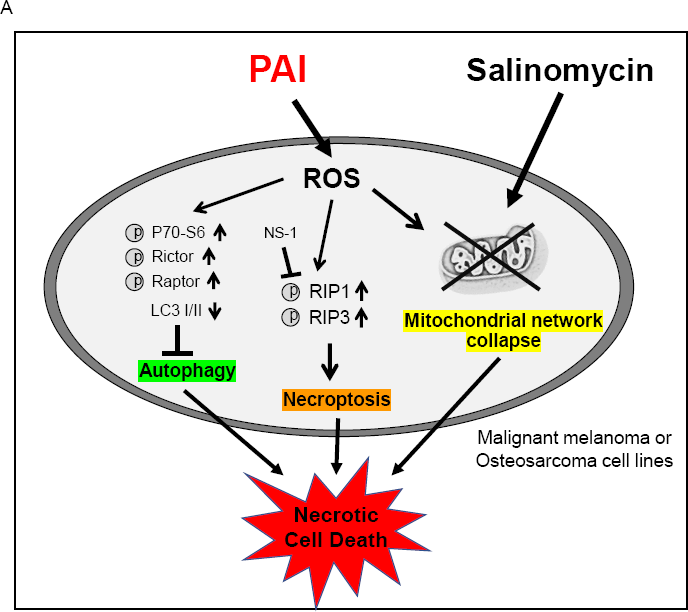

Both direct NTAP irradiation and PAL treatment have strengths and weaknesses. Direct plasma treatment is easy to control and chemically defines the active species. However, it is limited to cancers at the body surface because of its short outreach. PAL treatment is a rather indirect and more complicated approach compared to direct plasma irradiation. NTAP irradiation to liquids results in the production of various active chemical species, which are challenging to define. Nevertheless, PALs may be a promising approach for cancer treatment because they can be readily applied to all types of cancers, including leukemia and lymphomas, by endoscope and infusion. Although PAI induced cell death in MM and OS cells *in vitro* and significantly reduced OS tumor growth *in vivo*, the tumor could not be eliminated, demonstrating a limitation to PAI monotherapy. In clinical practice, it is difficult to treat cancer patients by mono chemotherapy. Considering this preservation of the residual tumor’s aggressiveness, the combination of PAI therapy with a chemotherapeutic drug may be a viable option as a new treatment regime, particularly if the combined drug itself is effective and safe. In the present study, we found that Sal is a candidate that meets these demands. As this compound can selectively kill cancer stem cells and multidrug-resistant cancer cells (Pellegrini et al., 2016; Yue et al., 2013; Klose et al., 2014), combination use of PAI and Sal may be useful for killing cancer stem cells and multidrug-resistant cancers that are tolerant to multidisciplinary treatment and cause tumor recurrence. Recent studies revealed that Sal causes significant weight loss in mice and toxicity to the male reproductive system and neuronal cells *in vivo* (Dewangan et al., 2017). Strikingly, combinatorial administration of Sal and other drugs such as gemcitabine, paclitaxel, and cisplatin ameliorate Sal toxicity show synergistic effects (Dewangan et al., 2017). Therefore, combination therapy with another drug rather than monotherapy is more promising and practical for the clinical use of Sal. In this regard, PAI and Sal mutually act as adjuvants to one another in killing OS cells (Figure 4). Thus, combined use of PAI and Sal can significantly reduce the windows of effective doses, thereby preventing adverse effects.

Understanding of the components of PAI remains challenging. Various types of analysis methods such as electron spin resonance and high-pressure liquid chromatography may be useful for addressing this issue. Notably, we observed that PAI was much more effective than H_2_O_2_ (100 μM), which was 2-fold higher than that contained in PAI (50%) (Figure 1). These observations suggest that another oxidant participates in the effect of PAI. Considering the chemical component of infusion fluid, PAI may be similar to plasma-activated PBS, which mainly contains H_2_O_2_, NO_2_^-^, and NO_3_^-^(4). H_2_O_2_ and NO_2_^-^ can synergistically induce cell death in human colorectal and MM cells. Therefore, nitrogen oxides may function as oxidants with a role in oxidative stress and cell death caused by PAI. Further studies are needed to evaluate these possibilities.

In conclusion, combination administration of PAI and Sal may be a safe and effective approach for treating apoptosis-resistant cancers such as MM and OS.

## 6. AUTHOR CONTRIBUTIONS

T.A. and Y.S.-K. designed the experiments and supervised the completion of this work. T.A., J.I. H.H. and Y.S.-K. verified the analytical methods. T.A., T.O., Y.Y., and Y.S.-K. contributed to preparation of resources. Ma.S.K. (Manami Suzuki-Karasaki), T.A. and Y.S.-K. performed the analytic calculations. Ma.S.K. (Manami Suzuki-Karasaki), T.A., Mi.S.K. (Miki Suzuki-Karasaki), and Y.S.-K. performed the experiments. T.A., H.H., Ma.S.K. (Manami Suzuki-Karasaki), and Y.S.-K. analyzed and validated the results. Ma.S.K. (Manami Suzuki-Karasaki), T.A., T.O., Y.Y., and Y.S.-K. wrote the original draft. T.A. and Y.S.-K. reviewed and edited the manuscript. T.A., T.O., Y.Y., H.H., and Y.S.-K. were responsible for funding acquisition. All authors discussed the results, contributed to and approved the final manuscript.

## 7. FUNDING

This research was funded by JSPS KAKENHI; grant numbers 15K09792, 17K10988, 18K08279, and 18K09121.

## 8. ACKNOWLEDGEMENTS

We thank the JCRB Cell Bank of National Institutes of Biomedical Innovation, Health, and Nutrition (Osaka, Japan) and Riken BioResource Center (Tsukuba, Japan) for providing cell lines. We thank M Ubagai and C Chino for their technical assistance. We also appreciate Dr. H Miyahara (Plasma Concept Tokyo) for invaluable suggestions in setting the plasma experiments.

## 9. Contribution to the field statement (193 words)

Osteosarcomas (OS) are malignant bone tumors. Advances in chemotherapy and surgery have increased the 5-year survival rate for patients. However, no other new drug with a high response rate has been developed and it remains difficult to improve prognosis using conventional chemotherapeutic regimens. Therefore, an innovative therapy for treating OS is urgently required. Here, we propose that combined administration of PAI and salinomycin (Sal) is a novel approach for treating OS. PAI is a new anti-cancer agent that is made by irradiating non-thermal atmospheric plasma to an infusion fluid, while Sal is an emerging anticancer cell agent. Either agent killed OS cell lines and mutually operated as adjuvants *in vitro*. Combined administration of PAI and Sal was much more effective than single agent application in reducing the growth and lung metastasis of OS allografts with minimal adverse effects. PAI explicitly induced necroptosis via the receptor-interacting protein 1/3 pathway, while suppressed the ambient autophagic flux by activating the mammalian target of rapamycin pathway. On the other hand, Sal promoted autophagy. Moreover, Sal exacerbated the mitochondrial network collapse caused by PAI. Thus, PAI and Sal seems to act synergistically by targeting autophagy and mitochondrial dynamics.

## 10. CONFLICTS OF INTEREST

Ma.S.K. (Manami Suzuki-Karasaki), Mi.S.K. (Miki Suzuki-Karasaki), and Y.S.K. are employees of the Non-Profit Institute Plasma ChemiBio Laboratory. The remaining authors have no conflicts of interest. The funders had no role in the design of the study; in the collection, analyses, or interpretation of data; in the writing of the manuscript, or in the decision to publish the results.

---

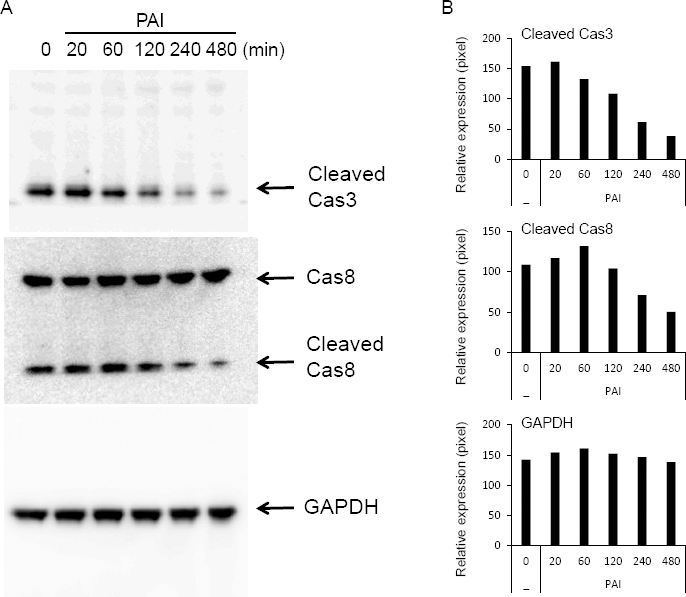

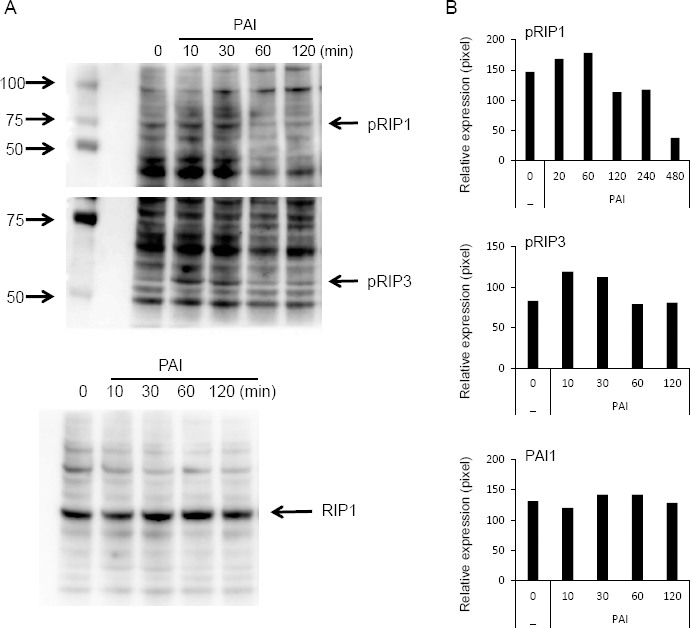

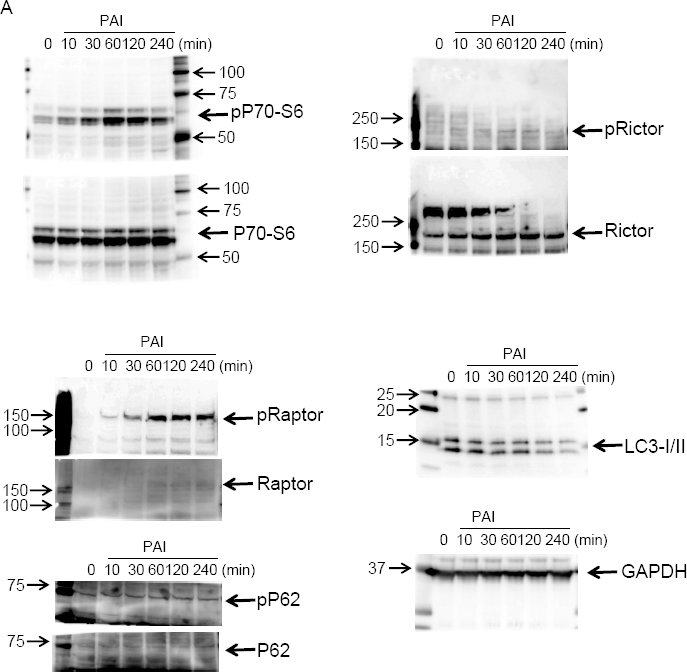

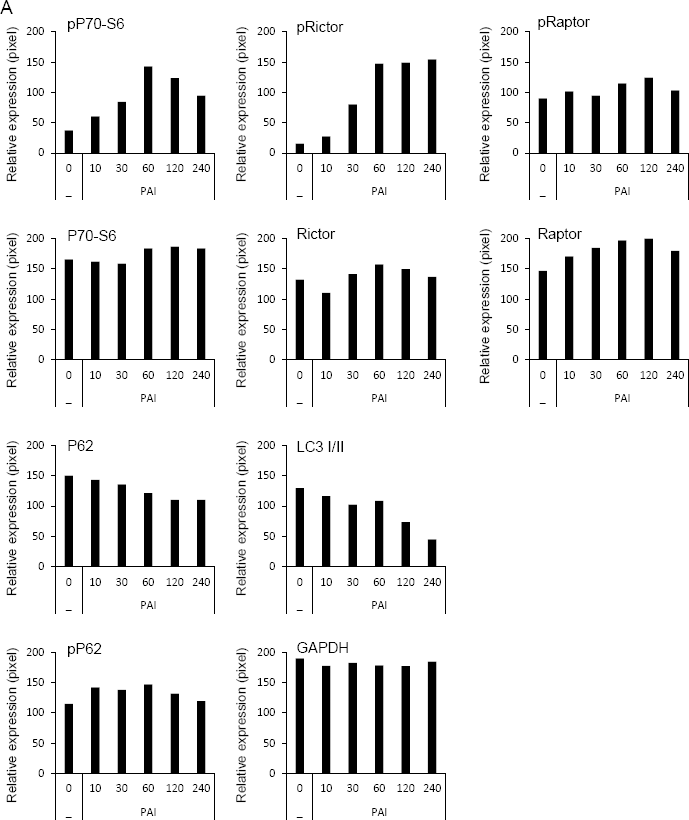

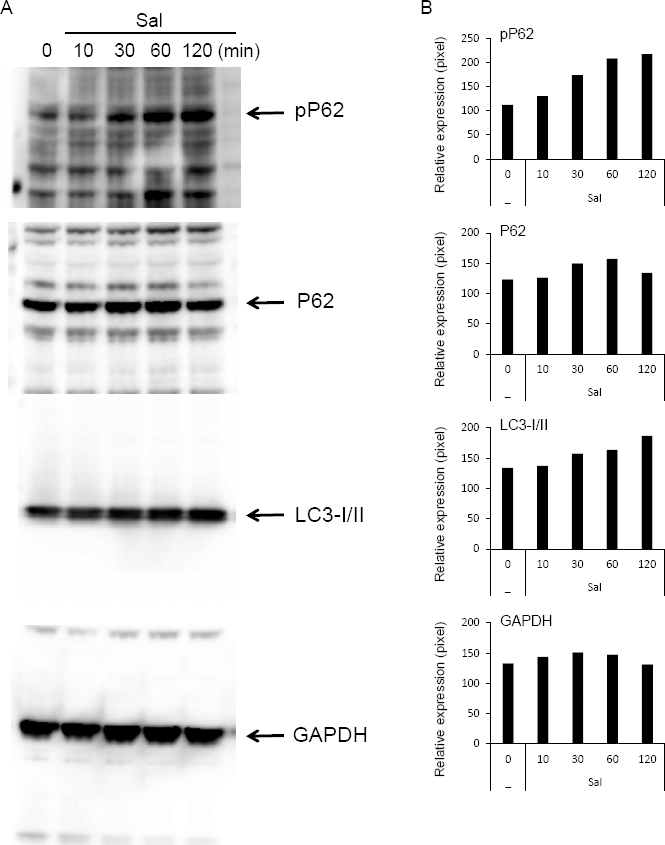

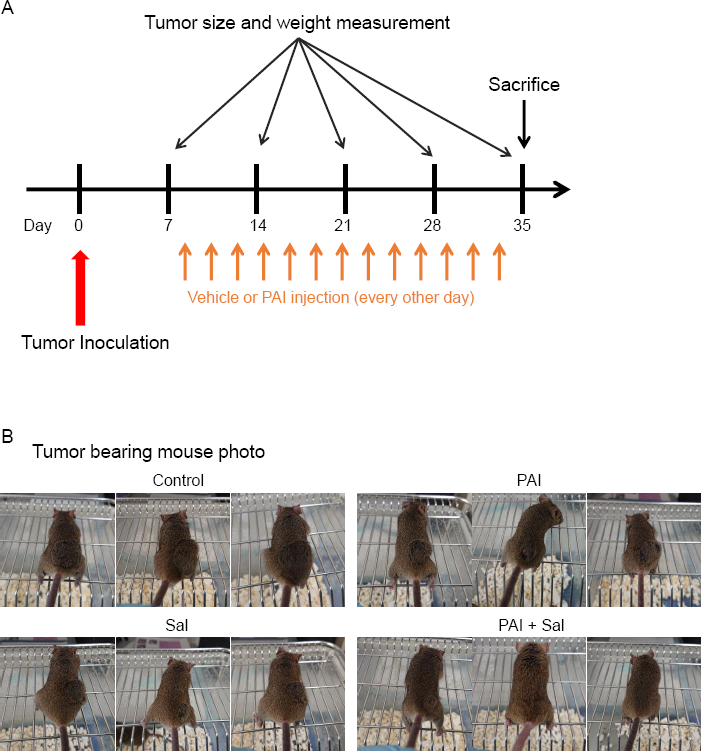

