## Supplementary material for "Synergistic anticancer effect of plasma-activated infusion and salinomycin by targeting autophagy and mitochondrial morphology": Supplemenal

### Supplementary Figure 1

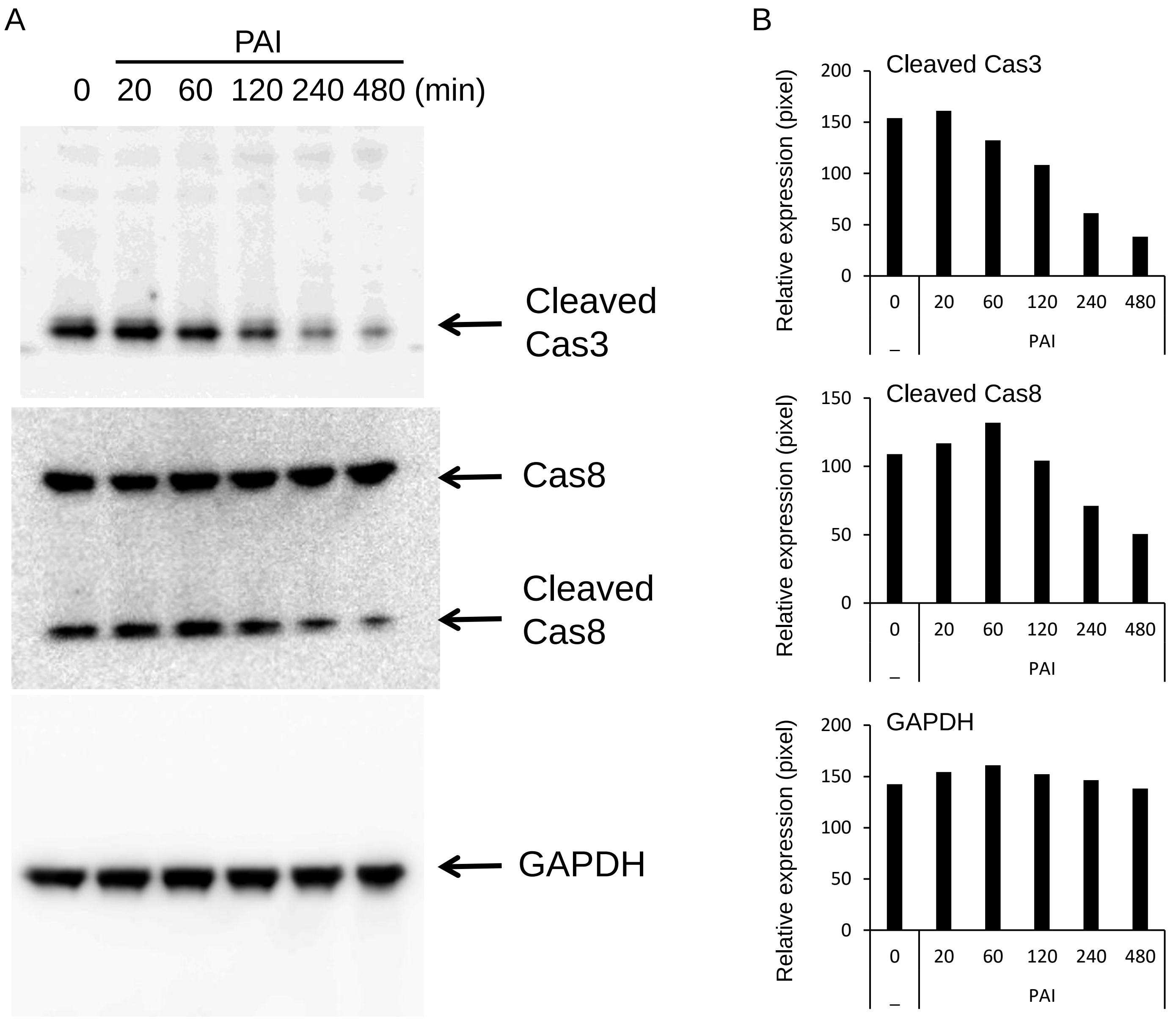

### Supplementary Figure 2

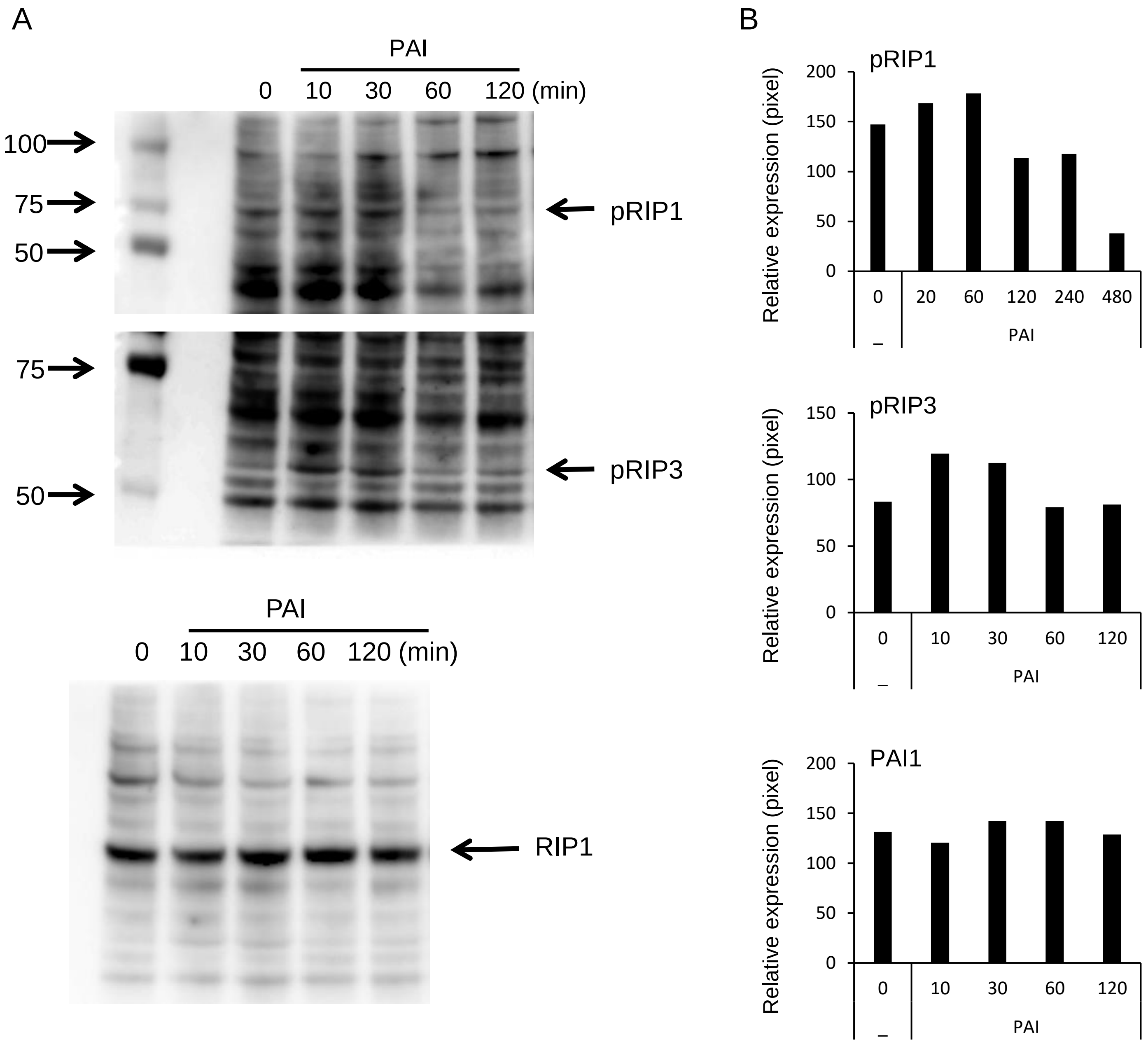

### Supplementary Figure 3

A

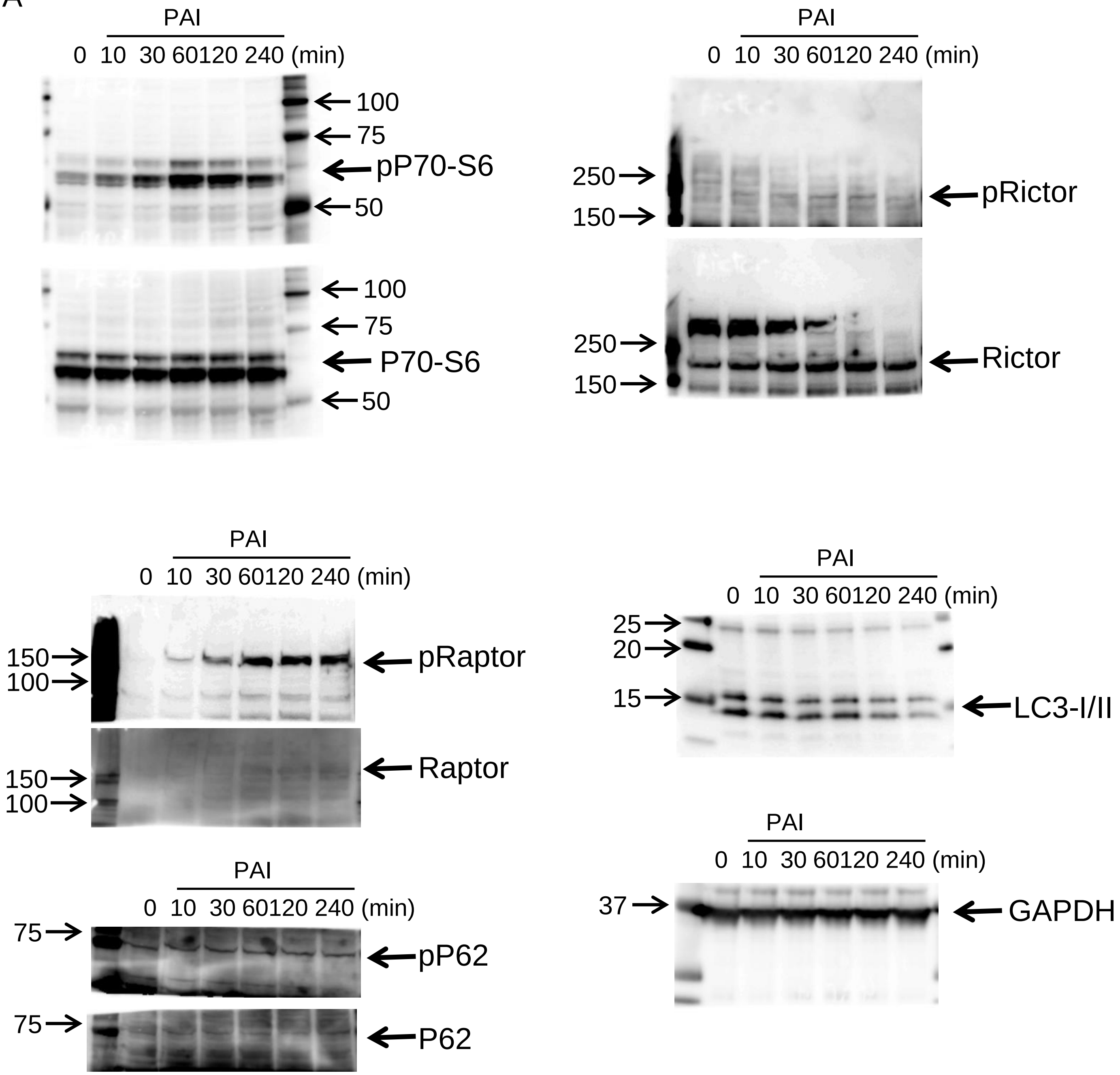

### Supplementary Figure 4

A

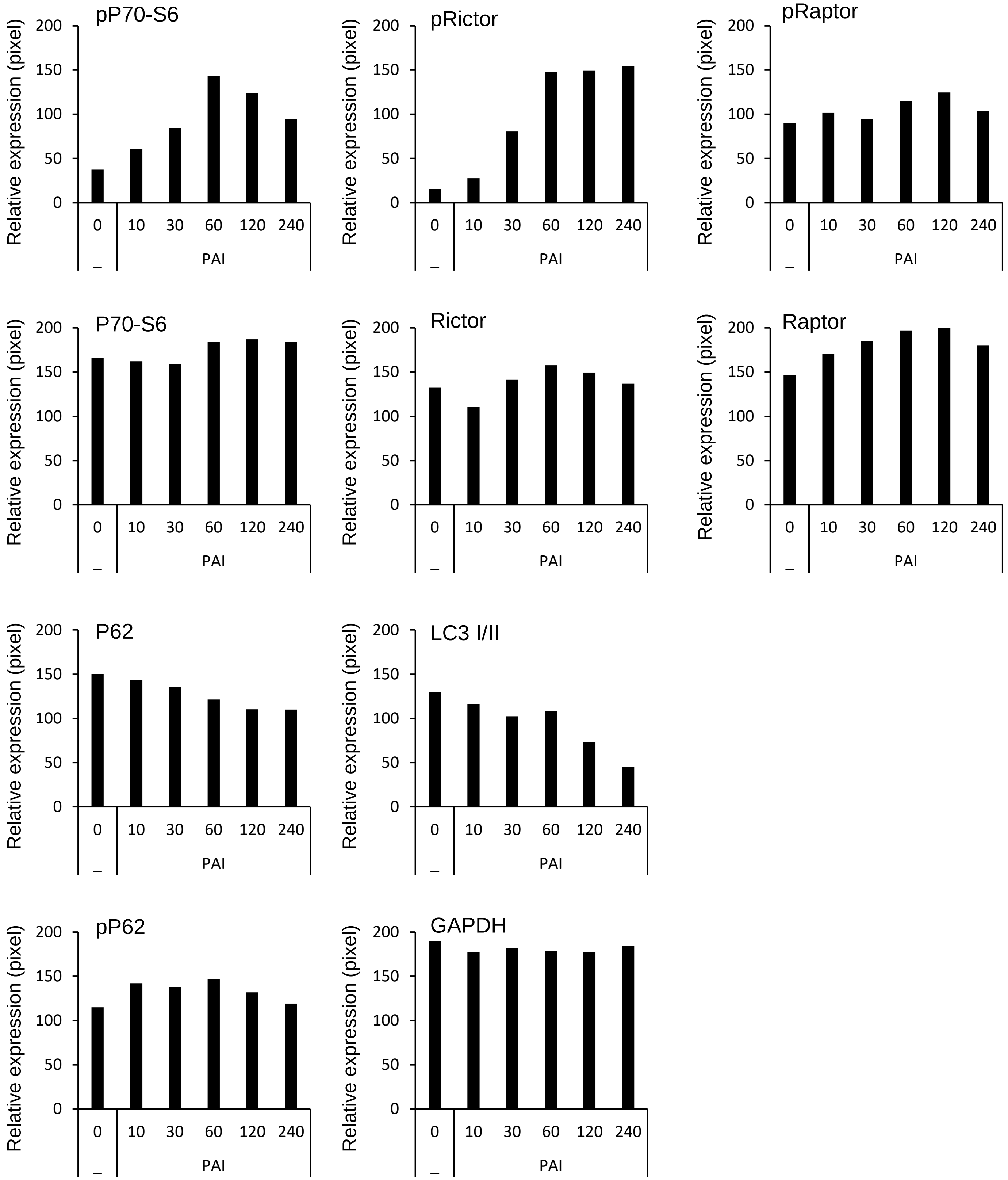

### Supplementary Figure 5

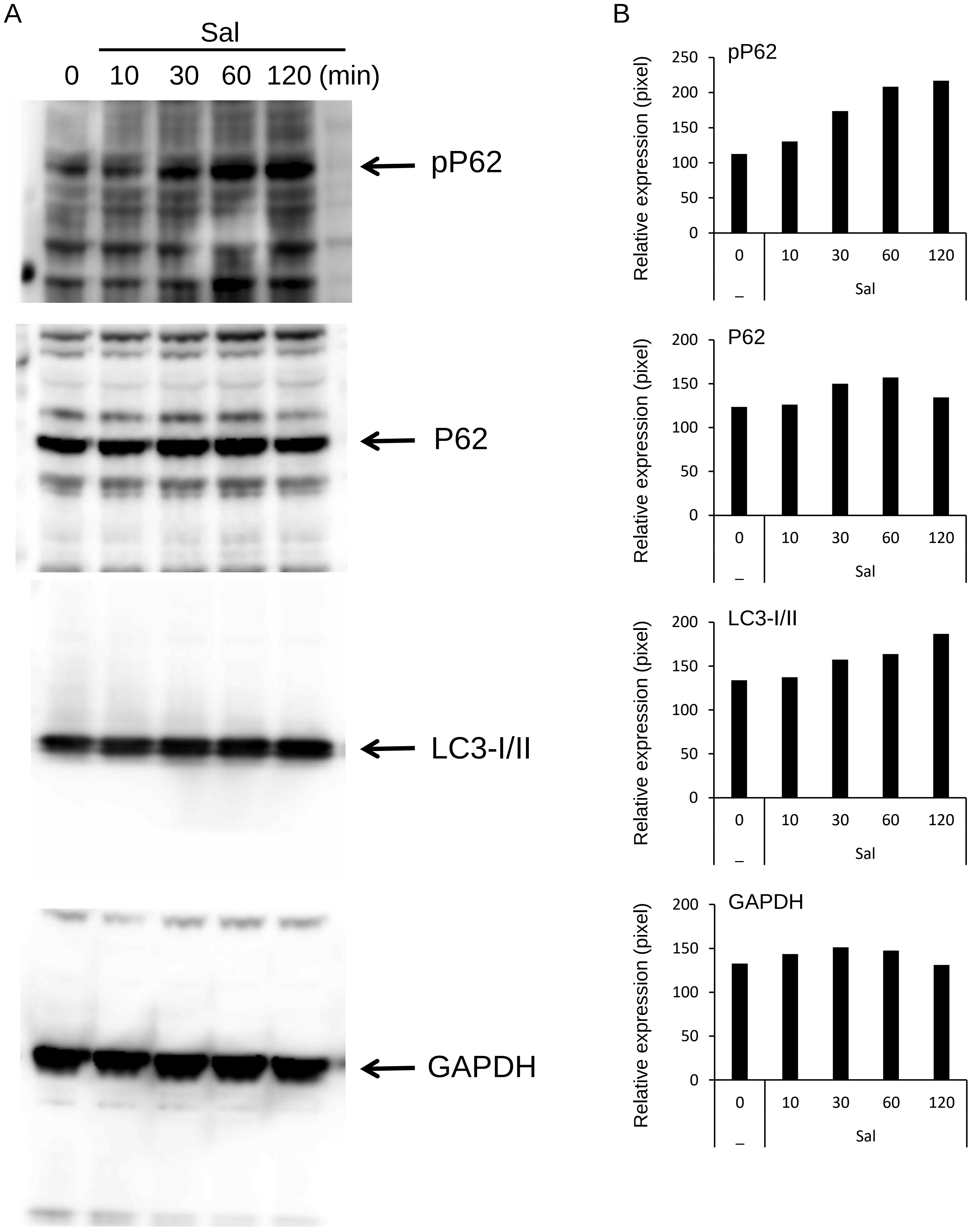

### Supplementary Figure 6

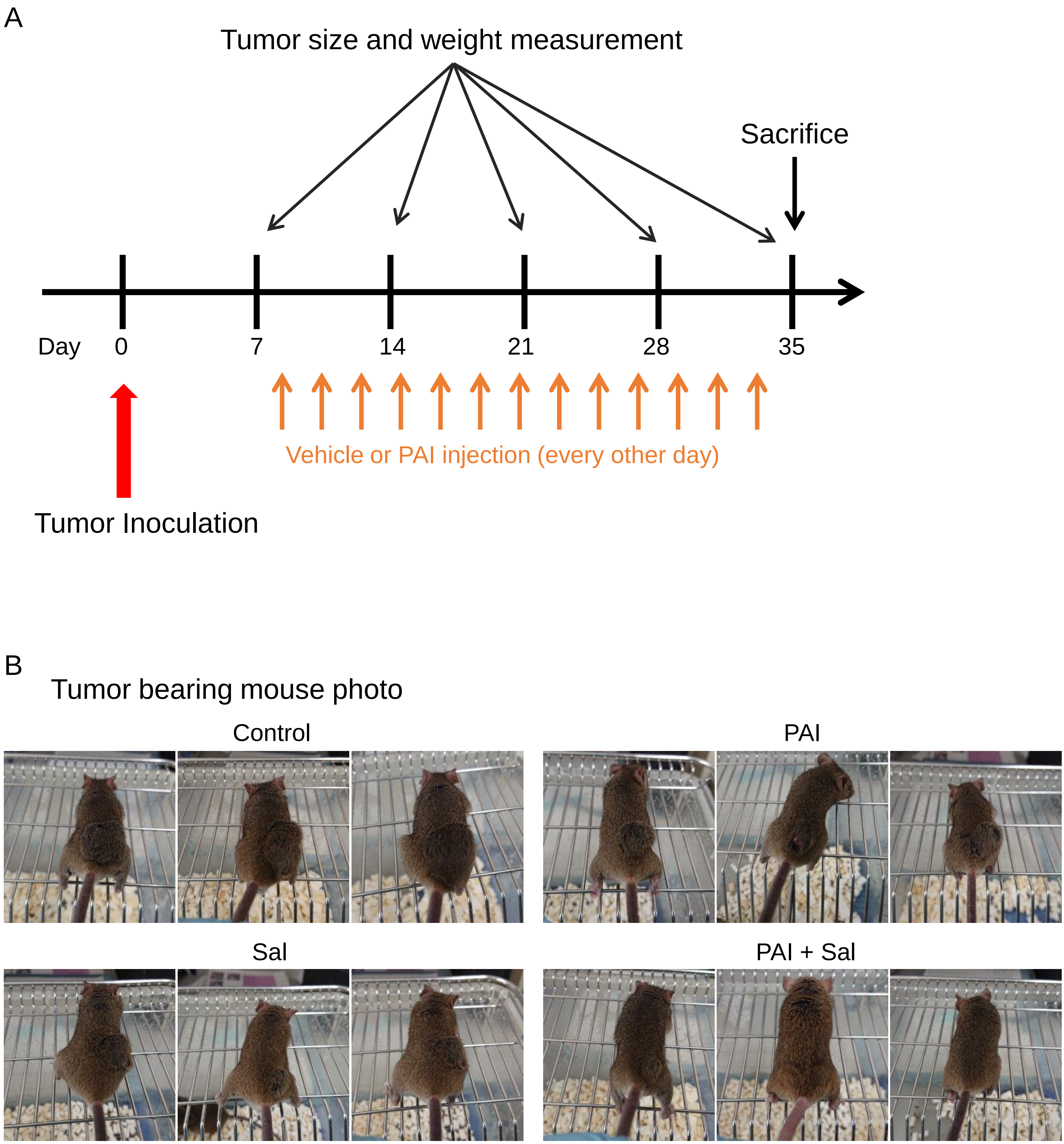
